# An integrative computational approach for obstacle avoidance during action selection

**DOI:** 10.1101/2025.11.17.688790

**Authors:** Shan Zhong, Nader Pouratian, Paul Schrater, Vassilios Christopoulos

**Affiliations:** Alfred E.Mann Department of Biomedical Engineering, University of Southern California, Los Angeles, CA, USA; Neuroscience graduate program, University of California Riverside, Riverside, CA, USA; Department of Neurological Surgery, UT Southwestern Medical Center, Dallas, TX, USA; Department of Computer Science and Engineering, University of Minnesota, Minneapolis, MN, USA; Department of Bioengineering, University of California Riverside, Riverside, CA, USA; Department of Neurological Surgery, Keck School of Medicine, University of Southern California, Los Angeles, CA, USA; Neurorestoration Center, Keck School of Medicine, University of Southern California, Los Angeles, CA, USA

## Abstract

Action selection in cluttered environments, where individuals must simultaneously pursue goals and avoid obstacles, presents a significant challenge for the brain. To understand the underlying mechanisms of action selection in such contexts, we propose a computational model that extends stochastic optimal control theory through a novel framework for obstacle avoidance. The model decomposes action selection as a weighted combination of individual control policies, each generated for either target approach or obstacle avoidance. By integrating value information from goals, obstacles, and actions into a unified measure of “relative desirability”, the model dynamically determines the contribution of each policy to the overall action selection process. We evaluated the framework using simulated target-reaching tasks in cluttered environments based on previous human studies. The results showed that the model captures key features of human motor behavior, including the influence of obstacle properties on movement trajectories and the transient tendency to initiate movements toward obstacles before avoidance. This work offers new insights into the dynamic interaction between approach and avoidance behaviors, providing a comprehensive framework for understanding action selection in complex and naturalistic settings.

**Author Summary:** Every day, we make countless movements that require us to reach for something while avoiding obstacles—for example, grabbing a cup without knocking over nearby objects. How the brain selects such actions in cluttered environments remains poorly understood. In this study, we developed a computational model that explains how people plan and control movements when both goals and obstacles are present. Our model assumes that the brain prepares multiple potential actions at once—some for reaching the goal and others for avoiding obstacles—and then combines them based on how desirable each option is at any moment. Using computer simulations of reaching movements, we found that the model reproduced key patterns observed in human behavior, including the brief tendency to move toward an obstacle before steering away from it. By showing how the brain might continuously balance approach and avoidance drives, our work offers a new way to understand how people make rapid and flexible movement decisions in complex, everyday settings.

## 1 Introduction

Consider a hypothetical scenario in which you are invited to a formal dinner and seated at a table with a hot bowl of soup in front of you, a glass of wine just behind it, a salt shaker in the center, and various other items nearby. Each of these objects can act as either a goal or an obstacle, depending on your intentions. For instance, if you want to take a sip of wine, the glass becomes your target. But if someone asks you to pass the salt shaker, the same glass now turns into a potential obstacle. Accidentally knocking it over could create an awkward or even embarrassing situation. The complexity of such interactions increases when certain actions can lead to positive or negative outcomes. Despite the importance of these decisions in everyday life, the mechanisms that allow the brain to select actions that achieve goals while avoiding obstacles remain poorly understood.

Research has shown that goal-directed reaching is not planned in isolation but is continuously shaped by the surrounding visual environment [1, 2]. This influence is particularly evident in tasks involving multiple potential targets, where initial reach trajectories are often biased toward a spatial average of the possible goal locations - a phenomenon known as *spatial averaging* [3–5]. Similarly, the presence of non-target objects systematically alters the path of the hand, causing trajectories to deviate even when the objects sometimes do not directly obstruct the most efficient route to the goal [6, 7].

One theoretical framework that accounts for these dynamic interactions is the “affordance competition hypothesis”, which posits that the brain continuously specifies and prepares multiple potential actions afforded by the environment [8, 9]. These partially prepared motor plans then compete for selection in a dynamic process that unfolds over time [10]. This theory is supported by neurophysiological evidence demonstrating the parallel encoding of multiple potential movements in sensorimotor regions [11–14], as well as findings showing that preparing for several possible actions increases motor variability [15]. Within this framework, reaching for a target while avoiding an obstacle can be conceptualized as a competition between a plan to approach the goal and one or more plans to avoid the obstacles.

A key challenge within the affordance competition hypothesis lies in specifying the computational mechanisms that determine how competing motor plans are dynamically weighted and integrated in real-time. This process involves integrating heterogeneous sources of information — such as expected rewards, action costs, and collision risks —which are often expressed in different “currencies” [16, 17]. To address this, previous work has proposed converting these disparate signals into a common probabilistic currency that encodes the *relative desirability*, thereby guiding the selection among competing goals [5, 17, 18]. However, it remains unclear how this framework extends to more complex scenarios involving simultaneous approaching and avoidance behaviors. Early computational models of obstacle avoidance, such as those based on artificial potential fields (APFs) [19, 20], treated obstacles as sources of repulsion. Although conceptually elegant in its simplicity, the classical APF approach suffers from notable limitations, most prominently the tendency to get trapped in local minima [21, 22]. In robotics, these challenges have motivated both the refinements to the potential field formulation [23, 24] and the emergence of more sophisticated techniques broadly classified as heuristic, predictive, or learning-based approaches [25]. Reinforcement Learning (RL), for instance, enables agents to learn optimal obstacle avoidance policies through trial-and-error iterations with their environment, offering enhanced adaptability to complex and dynamic contexts [26, 27]. Other modern techniques, including predictive methods such as model predictive control (MPC) and various swarm-based optimization algorithms, further expand the ability to solve obstacle avoidance problems that traditional methods struggle to manage [25]. While these advanced methods provide powerful solutions for autonomous navigation, their primary objective is often optimal performance in robotic systems, not necessarily biological plausibility. The goal of modeling human motor control is different; it seeks to understand the specific computational principles that give rise to observed human behaviors. For instance, the tendency for reaching trajectories to sometimes initially deviate towards a non-target object before curving away suggests a more complex integration of competing motor plans than simple repulsion or pure optimization would predict [2].

In the current study, we take a step towards a unified computational theory for action selection by extending the probabilistic framework introduced by Christopoulos et al. [17] - originally developed for multi-target choice - to the specific problem of reaching in cluttered environments. In our model, the final motor command is formulated as a dynamically weighted mixture of policies for target approach and obstacle avoidance, with each policy generated by a stochastic optimal controller [28]. We show this model captures key features of human reaching behavior, including the characteristic initial deviation of trajectories toward an obstacle. More broadly, this framework provides a principled means to describe the continuous competition between approach and avoidance drives, opening new avenues to investigate the neural computations that govern goal-directed movements in complex and cluttered environments.

## 2 Results

### 2.1 A computational model for selecting action plans in complex environments

The architecture of the model is a series of optimal control schemes associated with different targets/obstacles, as is shown in Fig.1. In stochastic optimal control theory, the objective is to determine a control policy that minimizes a cost function representing the trade-off between effort and goal attainment. Each optimal control scheme generates a specific action plan (policy) 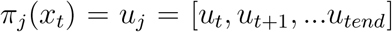, which defines a sequence of motor action plans *u* starting from the current state *x*_*t*_ and directed towards the desired goal *j*. The performance of a policy is evaluated by a cost function 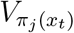, which computes the expected cumulative cost (i.e., effort) required to reach goal *j* from the current state while minimizing deviation from the desired final state.

**Figure 1.**
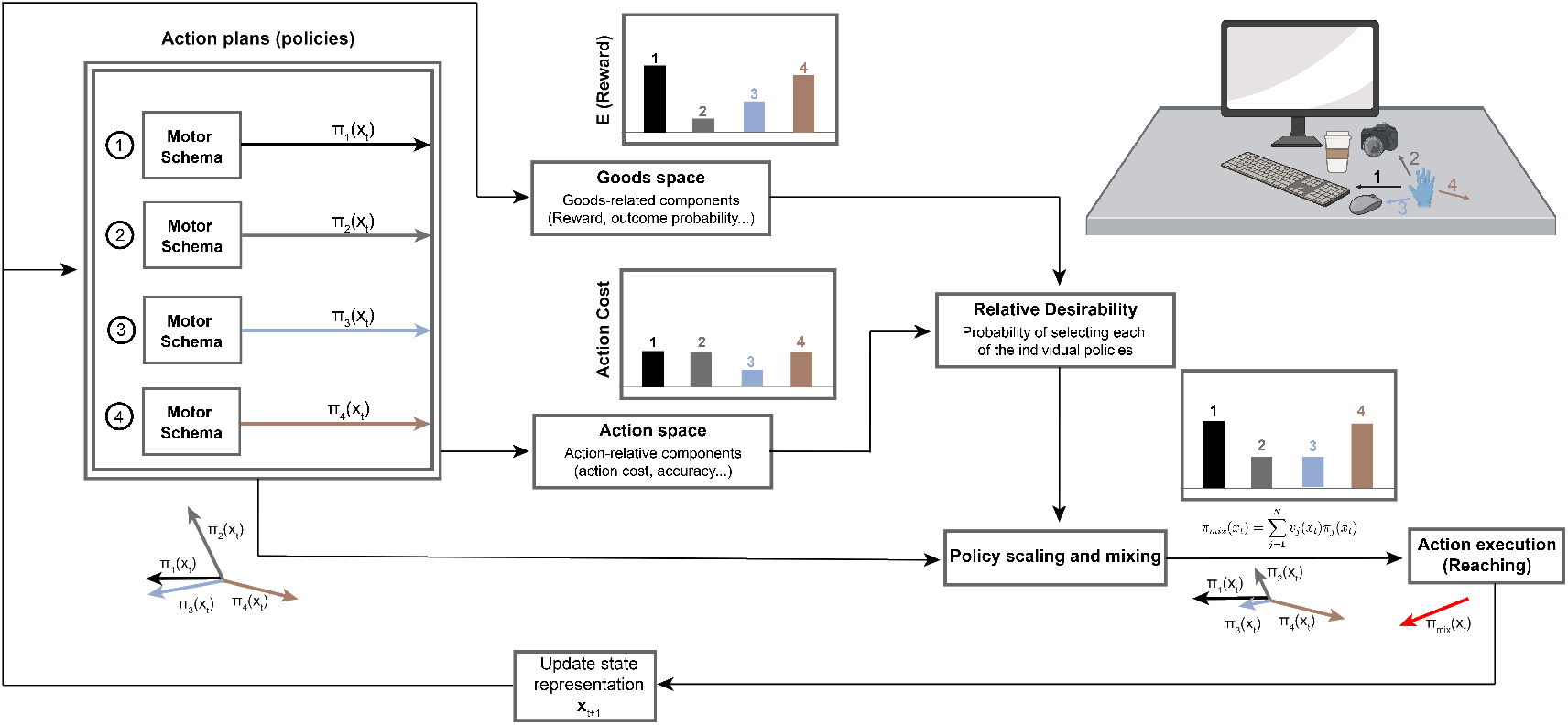
Model Architecture. The model consists of a series of optimal control schemes associated with different targets/obstacles. In this hypothetical scenario, a hand is positioned over a desk with different objects, among which three of them are potential targets (a keyboard, a computer mouse, and a camera), and one is a stationary obstacle (a coffee cup), which are located at different distances from the hand’s current state *x*_*t*_. Under these conditions, the motor schemas associated with each of these four objects are activated and generate four action plans *π*_1_ to *π*_4_. Note that *π*_4_ is negated since it corresponds to the obstacle. At each time t, the desirabilities of each of these action plans are computed based on their action-related components and goods-related components. Then these two values are combined to make an overall relative desirability. The final policy that the hand executes is a weighted mixture of the individual policies weighted by the relative desirability values.

Consider the reaching scenario in Fig.1, where a hand (at state *x*_*t*_) is positioned over a desk containing different target objects (a keyboard, a mouse, and a camera) located at distinct positions and distances, along with a stationary obstacle (a coffee cup) that must be avoided during reaching movements. Under these conditions, the optimal control schemes associated with the targets and obstacle are simultaneously activated, generating four action plans (*π*_1_ to *π*_4_) to move towards the targets and obstacle. The action plan directed toward the obstacle is inverted (negated) to approximate the optimal policy for avoiding it. Each action plan is associated with an action-related component, reflecting the effort required to execute that plan. For instance, it is typically less “costly” for the hand to move directly towards a target than to follow a curved trajectory. The targets are also associated with goods-related components, such as expected utility and probability of successful acquisition. For instance, reaching for the keyboard (the primary task object) offers higher utility than reaching for the mouse (a secondary object) in the current scenario. Furthermore, the obstacle is associated with obstacle-related components, such as the potential cost incurred upon collision (e.g., spilling coffee) and the probability of such a collision occurring based on the hand’s trajectory and the cup’s proximity. Consequently, the hand exerts greater effort to maintain safe clearance from the coffee cup when its position creates a higher collision risk.

In this study, we simplified the model by defining the goal-related term based on the probability of obstacle collision, as it is challenging to explicitly quantify the trade-off between the reward for reaching the target and the penalty for colliding with obstacles (Fig.1). Consequently, the obstacle-related components are incorporated into the goods-related term. During the reach, the effort cost, the probabilities of reaching the target, and the probability of avoiding obstacle all change dynamically. The individual must therefore integrate information from disparate sources to compute the most advantageous action at each moment. To achieve this, we employ a probabilistic framework that integrates these value signals into a common currency, termed *relative desirability* value [17]. The relative desirability combines information from both the action and goods space, representing the probability of selecting each individual action plan. The resulting optimal policy is then expressed as a mixture of action plans, weighted by their relative desirability values.

### 2.2 Collision Avoidance Reshapes Relative Desirability Distributions

To better understand how action cost and collision probability influence reaching behavior during obstacle avoidance, we visualized the spatial distribution of relative desirability under two conditions: (1) an action cost-driven scenario, in which only the action-related component is considered; and (2) a comprehensive scenario, which incorporates both the action-related and the goods-related (collision avoidance) components. For computational tractability and clarity of visualization, we focused the simulations on scenarios with a single target and a single obstacle, though the framework could generalize to multiple targets and obstacles as described in the Methods section. In the action cost-driven scenario, following the approach described in [17], we projected the four-dimensional desirability map (position-velocity space) onto two dimensions by constraining all the trajectories to originate from the position (0,0) with zero initial velocity. We defined a circular sampling region centered at (30,30) - i.e., the target location, with a radius of 75% of the origin- to-target distance, and allowed the trajectories to arrive at 359 evenly distributed spatial positions along the circle. At each position, the velocity was set radially outward from the origin through that spatial position, and speed was matched to that of an unobstructed optimal reaching movement at 75% of completion. The obstacle was positioned at (15,15). From each of these 359 positions on the circle, we computed the optimal trajectory toward the target while accounting for the obstacle. To incorporate variability, ten trials were simulated for each starting point. Representative trajectories are illustrated as arrows in Fig.2. Next, we discretized the space into a grid and computed multiple action-cost measures for each state, including the cost-to-go for reaching the target and avoiding the obstacle, as well as the relative desirability values associated with both. These computations produced a comprehensive state-space map that captures target-directed reaching behavior and obstacle avoidance (Fig.2). The spatial distributions of cost and relative desirability were then visualized using heat maps and vector fields (Fig.2).

**Figure 2.**
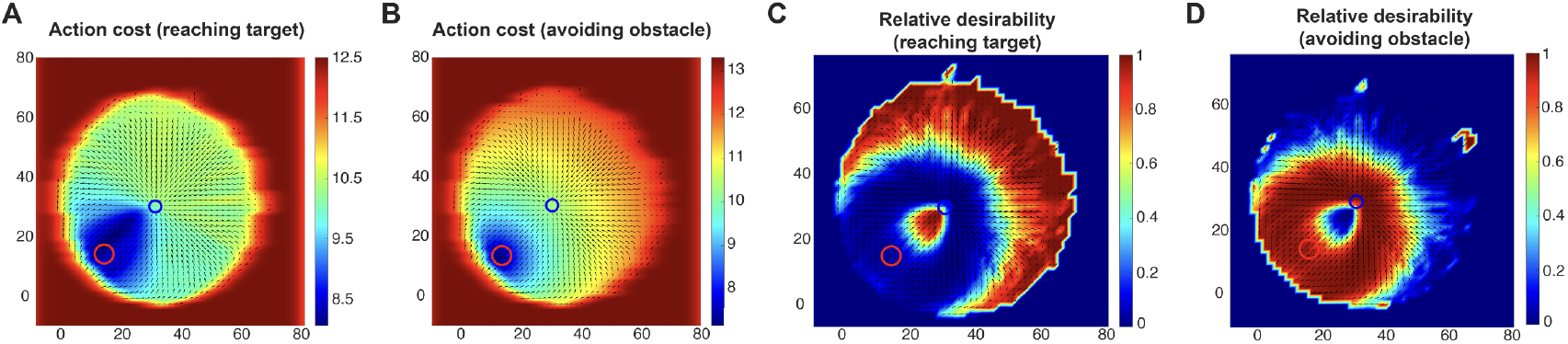
Action cost-driven relative desirability in obstacle avoidance. (A) Heat map of the action cost for reaching the target (blue circle) from different states. Warmer colors (reddish hues) represent higher-cost states, while cooler colors (blueish hues) represent lower-cost states. (B) Same as (A), but for avoiding the obstacle (red circle). (C) Heat map of the relative desirability for reaching the target. Warmer colors represent higher desirability, while cooler colors represent lower desirability. (D) Same as (C), but for avoiding the obstacle.

As shown in Fig.2A, the action cost for reaching the target is lowest in the bottom-left region, where minimal deviation in movement direction is required. In contrast, Fig.2B illustrates that the action cost associated with avoiding the obstacle is lowest in the vicinity of the obstacle. This counterintuitive outcome arises because the obstacle-avoidance policy is defined as the negation of the policy for reaching the obstacle, effectively creating an “anti-target” at the obstacle’s location. When the two action-cost maps are integrated to compute the relative desirability (Fig.2C and D), the resulting spatial distribution of policy weights appears more realistic. The desirability for reaching the target is high when the current state is distant from the obstacle (Fig.2C), consistent with intuitive expectations. However, the model still produces an unrealistic “rim” of high obstacle-avoidance desirability surrounding the target (Fig.2D). This occurs even when the trajectory is directed away from the obstacle and is close to the target, conditions under which avoidance behavior would be unlikely in natural settings. This discrepancy shows a limitation of relying solely on action cost, as it fails to account for the spatial relationship between the current movement direction and the position of the obstacle.

In the comprehensive scenario, we went beyond computing the cost of reaching the target and avoiding the obstacle, by also estimating the collision probability using an expanded obstacle representation. We introduced an expanded *collision zone* surrounding the actual obstacle, defined by two diameters: an inner diameter corresponding to the physical size (i.e., diameter) of the obstacle, and an outer diameter specifying the tolerance boundary within which a potential collision is still considered likely. Collision probability is computed based on the position of the hand relative to these boundaries. Points within the inner diameter are assigned 100% collision probability. For points between the inner and outer diameters, the probability decreases linearly from 100% to 0%, reflecting the graded nature of human perception and caution near obstacles (see Methods section for more details). Fig.3 applies the same visualization approach used in the action cost-driven scenario, but now incorporates the collision probability, resulting in a more comprehensive description of the obstacle avoidance task. In Fig.3A, we illustrate the collision probability, which reveals two distinct *collision cones*: one centered around the obstacle, as expected, and a second surrounding the target. The latter captures trajectories that, while aimed at the target, approach the obstacle dangerously closely. Within each cone, the collision probability peaks at the center and gradually decreases towards the periphery. By integrating these collision cones with the previously computed action costs presented in Fig.2, we observe remarkable changes in the relative desirability landscape (Fig.3B and C). Specifically, trajectories that approach or pass in close proximity to the obstacle are assigned lower desirability. The resulting maps provide a more accurate representation of how humans perceive and plan movements in cluttered environments with potential obstacles.

**Figure 3.**
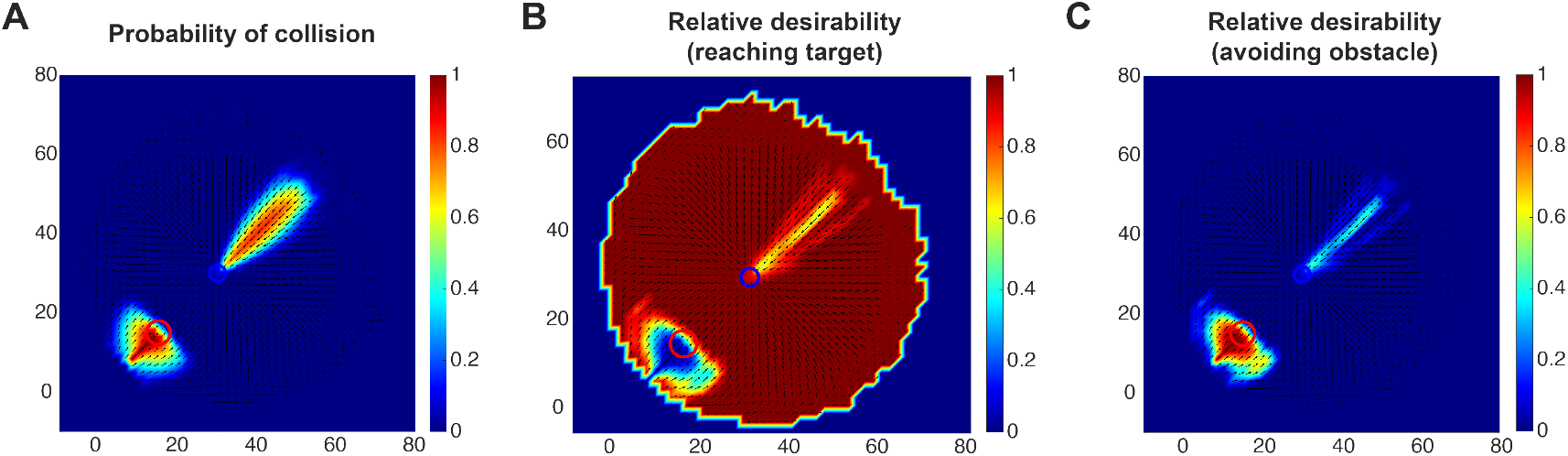
Comprehensive relative desirability in obstacle avoidance. (A) Heat map showing the probability of collision, where warmer colors denote higher probabilities and cooler colors represent lower ones. (B) Heat map of the relative desirability of reaching the target, taking into account both action costs and collision probability. Warmer colors represent higher desirability. (C) Same as (B), but illustrating the relative desirability associated with avoiding the obstacle.

### 2.3 Relative Desirability Predicts Reaching Behavior in Avoiding Obstacles

To validate the predictive capabilities of our model, we simulated reaching behaviors across multiple obstacle configurations. We focus on two key scenarios: one in which the obstacle lies directly in the path to the target, and another where the obstacle is nearby but does not obstruct the direct path. Fig.4 illustrates the first scenario, where an obstacle is positioned between the starting point and the target. To assess how the obstacle size influences behavior, we systematically increase the diameter of the obstacle across trials.

**Figure 4.**
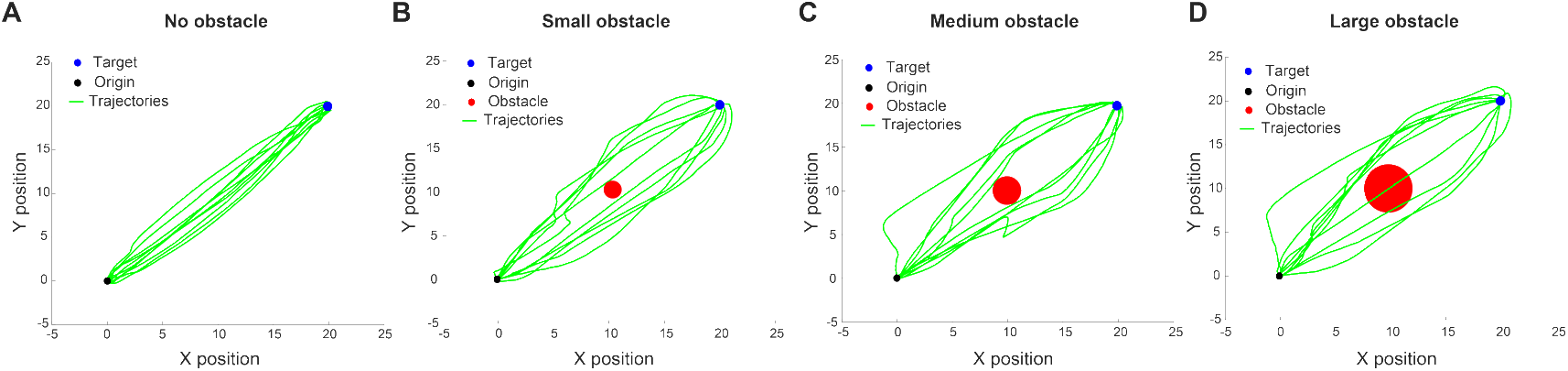
Simulated reaching trajectories with varying obstacle sizes along the direct path. Simulated reaching trajectories (green lines) for four conditions: (A) No obstacle, (B) small obstacle, (C) medium obstacle, and (D) large obstacle. The origin is indicated by a black dot, the target by a blue dot, and obstacles by red circles.

As shown in Fig.4, the model predicts distinct reaching behaviors across varying conditions. In the absence of an obstacle, trajectories to the target are direct and efficient. As an obstacle is introduced and its size progressively increases, the reaching movements begin to deviate, curving away from the obstacle’s location. For sufficiently large obstacles, the model occasionally predicts collisions, reflecting scenarios where the cost of avoidance outweighs the cost of collision. These predictions closely mirror natural human behavior, where larger obstacles tend to elicit wider avoidance paths and, in some cases, result in contact due to high avoidance cost.

Fig.5 illustrates the second scenario, in which the obstacle is positioned near - but not directly obstructing - the reaching path towards the target. We simulated 100 trials for each condition, with the obstacle placed either to the left (Fig.5A) or to the right (Fig.5B) of the direct path from the origin to the target. Interestingly, the simulations often produced trajectories that initially deviated towards the obstacle - an effect consistent with previous observations in human studies [1, 29]. In some cases, these trajectories subsequently curved around the obstacle before continuing towards the target. For visualization clarity, we separated these trajectories from the rest and computed their standard errors independently. In other instances, the trajectories deviated away from the obstacle even when it did not directly block the path, resulting in a rightward bias when the obstacle was on the left side (Fig.5A) and a leftward bias when it was on the right side (Fig.5B), consistent with findings from prior human studies [7,30]. These predictions capture the subtle influence of nearby obstacles on human reaching behavior, demonstrating that individuals tend to avoid the obstacle even when it does not directly interfere with the most efficient path to the target.

**Figure 5.**
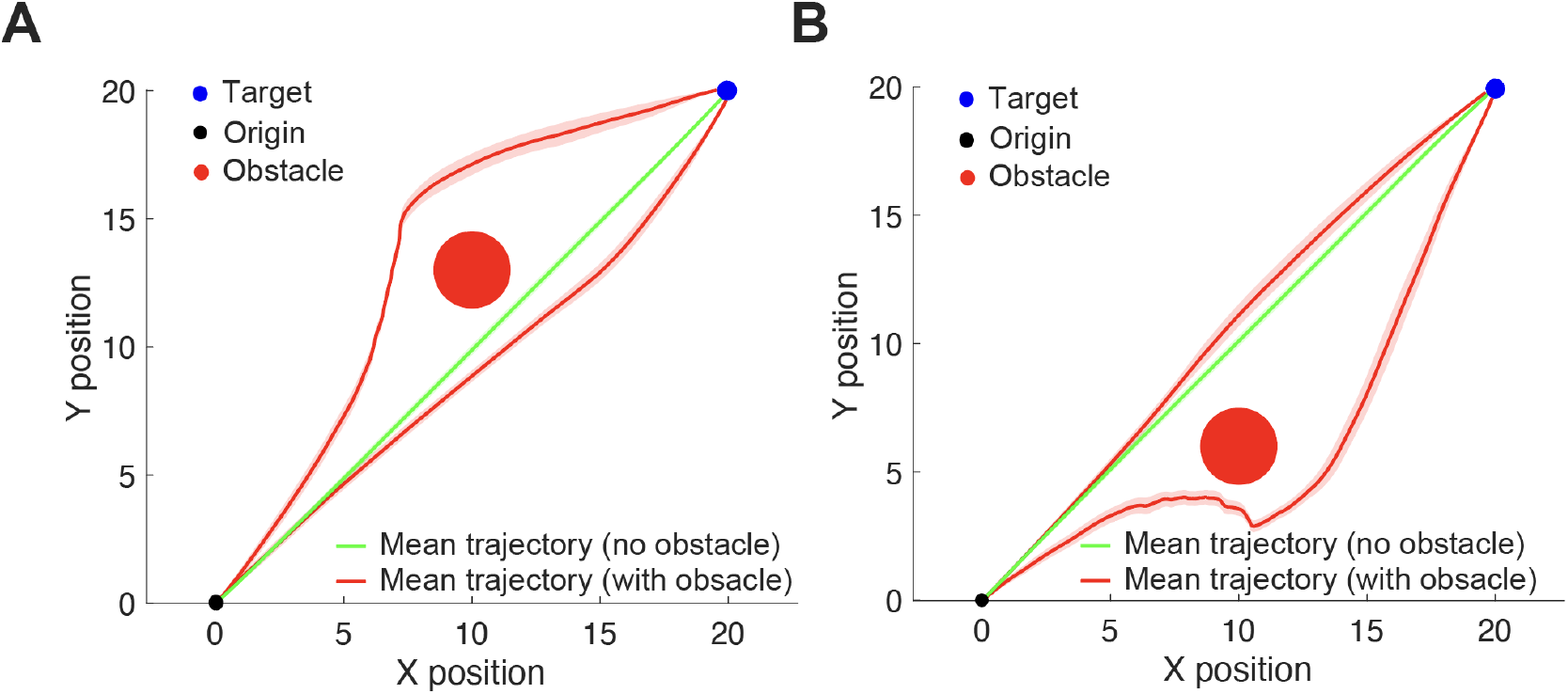
Simulated reaching trajectories with nearby obstacles. Mean simulated reaching trajectories with (red lines) and without (green lines) an obstacle (red circle) when reaching toward a target (blue circle). Shades represent standard error of the simulated trajectories. (A) Obstacle located to the left of the direct path. (B) Obstacle located to the right of the direct path.

Furthermore, we examined the performance of our model in more dynamic environments where the target location changes mid-flight. Fig.6 illustrates two such scenarios. In Fig.6A, the target jumps to a new position during the reaching movement, and we simulated two conditions: one with an obstacle (red circle) positioned along the direct path to the new target location, and one without any obstacle (green trajectories). In Fig.6B, we examined a similar target-jump scenario under three conditions: a no-obstacle (green trajectories), and two separate obstacle configurations (obstacle 1 - red circle and obstacle 2 - blue circle), in which an obstacle was positioned at different nearby locations that did not directly block the path to the new target location. Across all conditions, the model successfully generated trajectories that redirected toward the new target location. Importantly, trajectories deviated away from the obstacle, whether the obstacle directly obstructed the path (Fig.6A) or was merely nearby (Fig.6B). These simulations demonstrate that the model captures the flexible and adaptive nature of human motor control in dynamic environments, preserving obstacle-avoidance behavior when goals shift during movement execution.

**Figure 6.**
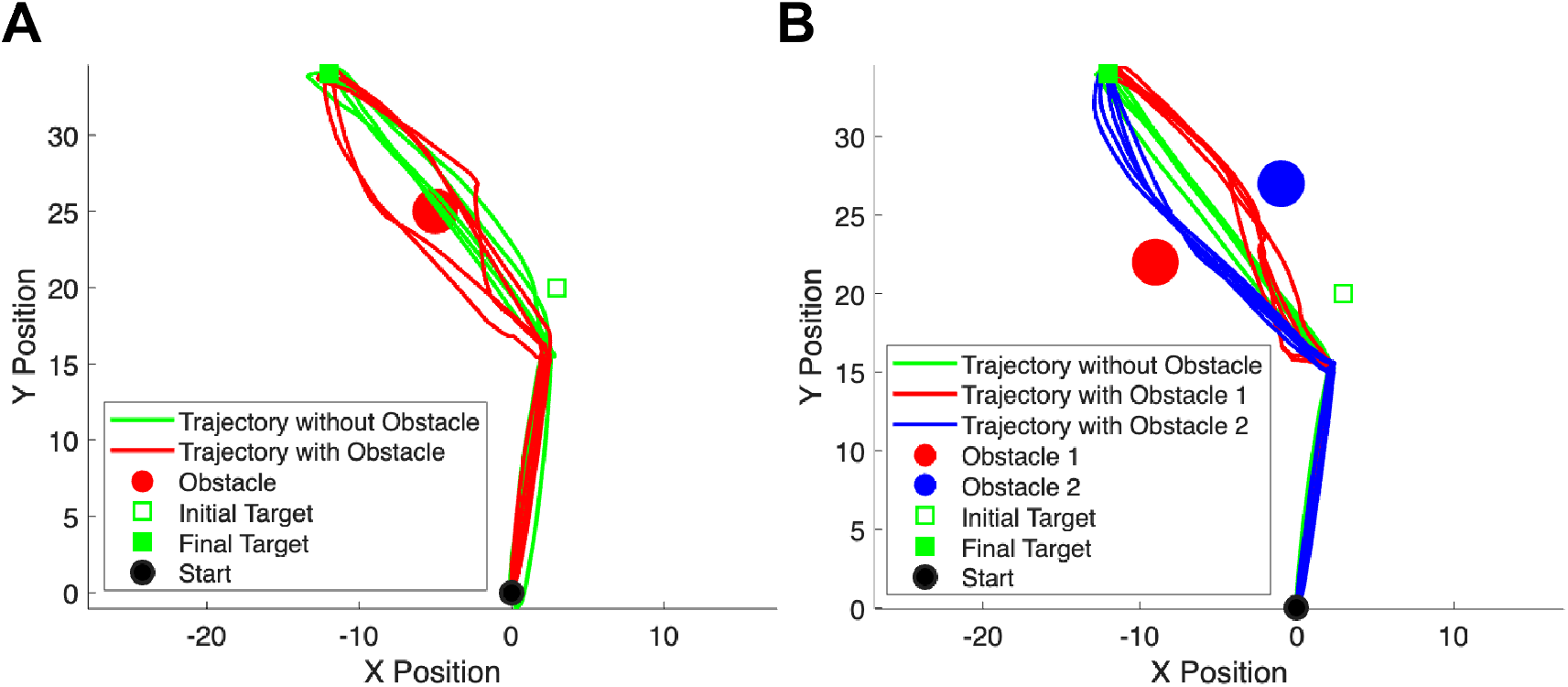
Simulated reaching trajectories in dynamic environments with mid-flight target changes. (A) Reaching behavior when the target jumps to a new location during movement, with (red) and without (green) an obstacle blocking the direct path to the final target. (B) Similar to (A), but with three conditions: no obstacle, and two different obstacle configurations (obstacle 1 or obstacle 2) positioned nearby but not directly blocking the path to the final target.

## 3 Discussion

### 3.1 General

The ability to navigate in cluttered environments while simultaneously approaching targets and avoiding obstacles is fundamental to everyday behavior, yet the underlying computational mechanisms remain poorly understood. Traditional computational approaches, such as APFs, treat obstacles as sources of simple repulsion but suffer from limitations like local minima trapping [21, 22]. While modern robotics has developed sophisticated solutions [25, 27], these approaches prioritize optimal performance over biological plausibility. The current study addresses this challenge by extending the probabilistic framework of Christopoulos et al. [17] from multi-target choice scenarios to the specific problem of reaching in cluttered environments. Building upon the affordance competition hypothesis [8, 9] and previous work on action-selection and value integration [17], we developed a model that integrates action-related costs, expected rewards, and potential losses associated with both targets and obstacles. The model decomposes the complex problem into a weighted mixture of individual control policies, each generated by a stochastic optimal control system to either move towards a target or avoid an obstacle. The relative desirability of each policy serves as a common currency for comparing and integrating the value information from disparate sources [17], consistent with the notion of a common neural currency for decision-making [31].

The results demonstrate that the model captures key aspects of human reaching behavior in the presence of both targets and obstacles, such as the tendency to deviate from the direct path to the target, as well as the occasional deviation towards the obstacle before curving around and turning to the target. Furthermore, the framework predicts a decrease in the desirability of movements toward a target when an obstacle is located immediately beyond it along the same movement direction—that is, when the obstacle does not impede the reach itself but is positioned just past the target. These results align with previous findings in human studies [3, 32, 33] and highlight the model’s ability to account for the spatial relationship between the current movement state and the obstacle. The ability of the model to reproduce these characteristic trajectory patterns, including initial deviations toward obstacles [2], demonstrates how competing motor plans can be dynamically integrated rather than simply competing through mutual inhibition.

### 3.2 Model Characteristics and Biological Plausibility

It is important to acknowledge that our model does not provide a computationally optimal solution to obstacle avoidance. The “policy negation” approach — i.e., computing the policy to reach an obstacle and then inverting it — serves as an approximation rather than the result of directly optimizing an avoidance cost function. This negation step is necessary because stochastic optimal control inherently requires a well-defined end goal, whereas obstacle avoidance lacks a specific endpoint to move toward. In contrast to target-directed movements, where the target location defines the optimal control goal, avoidance behavior must be inferred through the inversion of an approach policy. Thus, our objective is not to achieve mathematically optimal performance but to construct a biologically plausible model that captures key features of human motor behavior. Indeed, human movements exhibit substantial variability and deviations from optimality [34], reflecting the constraints of neural computation including uncertainty, processing delays, risk aversion, and so on. The value of our approach lies in its ability to reproduce realistic human-like behaviors, including seemingly inefficient patterns such as initial trajectory deviations towards obstacles [2].

By prioritizing biological plausibility over mathematical optimality, our model provides a framework for understanding how the brain might solve the obstacle avoidance problem in cluttered environments through integration of multiple motor plans, rather than optimizing a global cost function. Furthermore, although distinct from traditional obstacle avoidance algorithms, our model is not aimed to argue against the utility of these methods such as APF, RL or MPC for their intended applications in autonomous navigation and robotics [23, 25–27]. These algorithms are designed to achieve optimal performance, whereas our model is intended to provide an alternative and mechanistic insights into human motor control and to generate testable predictions about neural processes.

### 3.3 Limitations and future perspectives

By capturing key aspects of human obstacle avoidance behavior, the model also opens new avenues for refining and extending its underlying mechanisms. One such aspect is the treatment of collision probability, which is currently implemented through expanded collision zones. While this approach is computationally efficient and effective in reproducing qualitative behavioral patterns, it remains a simplified representation. Future approaches could incorporate probabilistic representations based on obstacle geometry, risk, and movement uncertainty [35]. Moreover, the risk aversion parameter *α* was treated as a fixed value in our simulations. However, evidence suggests substantial individual differences in risk-taking behavior [36, 37], showing that this parameter likely varies across subjects and contexts. Investigating how *α* relates to behavioral or neural markers could provide insights on the mechanisms underlying individual differences in risk-taking.

Furthermore, while the model successfully captures behavior in environments with static obstacles, extending it to more dynamic and complex contexts would offer a more rigorous assessment of its generalizability. Although the model architecture supports multiple targets and obstacles concurrently (as shown in Fig.1), the current simulations were limited to single target-obstacle pairs to characterize the fundamental behaviors. Future work should investigate how the model performs in more complex scenarios with multiple targets and obstacles, where the dynamic weighting of policies becomes more crucial. To guide and validate this extension, systematic comparisons between model predictions and human reaching behavior across various obstacle configurations, movement speeds and task constraints will be essential to testing the model’s predictive accuracy and refining its parameters. Such behavioral validation will also pave the way for tackling more complex real-world conditions involving multiple moving obstacles and goals, continuous goal adjustments, and predictive planning, which require a framework capable of integrating desirability principles within a dynamic architecture. Recent work has shown that the relative desirability framework, when combined with dynamic neural field (DNF) theory, could successfully and effectively model both the underlying behavioral and neural mechanisms of action selection, stopping and switching of actions [18, 38], suggesting that our principles of desirability and dynamic policy integration have the potential to naturally extend to cluttered, dynamic environments. Finally, exploring the neural substrates of obstacle avoidance represents a promising direction for future research. Electrophysiological recordings in premotor, motor, and parietal cortices, as well as in subcortical regions such as the subthalamic nucleus (STN), which is involved in action reprogramming and movement pausing, could provide valuable insights into the predictions of the proposed model.

## 3.4 Conclusion

In conclusion, the current study presents a biologically-plausible computational model that extends probabilistic action selection principles to obstacle avoidance scenarios. By dynamically updating the relative desirability of individual control policies, the model provides insights into how the brain may select actions in the face of both goals and potential obstacles. Future studies on the neural mechanisms of approach–avoidance behavior, informed by our computational framework, will be vital for elucidating how the brain achieves flexible and adaptive control in complex settings.

## 4 Methods

### 4.1 Model architecture

The model is comprised of a series of stochastic optimal control systems, each of them attached to an individual goal or an obstacle (Fig.1). Each controller *j* generates an optimal policy *π*_*j*_, which is a mapping between a current state and optimal actions, and an action cost function 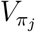 that describes the cost that is expected to accumulate following the policy *π*_*j*_ from the given state. To avoid any confusion, it is important to point out that a policy *π*_*j*_ is not a particular sequence of actions-it is a function that presents the best action plan (i.e., a sequence of actions ***u***_***j***_) to take from any state *x*_*t*_ to either acquire the goal *j* or to avoid the obstacle *i*-depending on whether the controller is attached to a goal or an obstacle - (i.e., 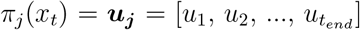).

The operations of the present theory can be easily understood in the context of a realistic action-selection scenario in an environment with many competing opportunities and risks. Consider the reaching scenario illustrated in Fig.1. A hand rests on a desk surrounded by three potential target objects—a keyboard, a mouse, and a camera—each located at different distances from the current hand position. The environment is cluttered and contains a stationary obstacle (a coffee cup) that must be avoided while reaching toward any of the targets. In such a situation, the control systems related to the goals and the obstacle become active and generate policies *π*_*i*_(*x*_*t*_), *i* = [1,4] to reach the targets and to avoid the obstacle. Each of these policies is accompanied by rewards or losses and action costs, and the value of them may change with time and previous actions. The reward values refer to the expected benefits for reaching each of the alternative targets on the desk. The loss values represent the expected cost associated with failing to avoid an obstacle (i.e., colliding with the coffee cup). Finally, the action cost refers to the cost (i.e., effort) required to implement each policy at a given state. Selecting between alternative policies requires integrating multiple decision variables online, that is, continuously and dynamically while acting. However, these decision variables are expressed in different “currencies”, making their integration into a unified value representation challenging. To address this, we adopt a probabilistic approach to integrate value information into a common currency. At first, the decision variables are compared within three distinct spaces: i) **goods-space**, where the expected reward values associated with individual goals are compared, ii) **obstacle-space**, where the expected loss values resulting from potential collisions are assessed—this space was simplified and integrated into the goods- space in the present study, iii) **action-space**, where the action costs of executing policies to achieve the goals and avoid obstacles are compared. The outputs from these comparisons are integrated into a scalar value named “relative desirability” and characterize how desirable it is to follow a particular policy with respect to the alternatives at a given time. It takes values between 0 (undesirable) and 1 (highly desirable) and is used to weight the contribution of the individual policies to the mixed-policy that will be followed by the hand at a given state.

### 4.2 Modeling action selection in dynamic environments

This section briefly presents the computational theory developed to model action-selection tasks in dynamic environments. More details about the model are presented analytically in [17]. We used a reaching simulated experiment as a paradigm, although the framework can be extended to model other tasks, such as grasping.

#### 4.2.1 Multiple competing goals in obstacle-free environments

The computational model is based on stochastic optimal control theory, which has been extensively used over the past years to model goal-directed movements, such as reaching to touch targets, pointing to a direction and grasping objects [28, 39, 40]. The basic idea of the optimal control theory is that movement policies *π*(*x*_*t*_), at any given state *x*_*t*_, are generated by optimizing performance criteria that include both goal- and action-related costs. We have proposed a probabilistic theory [17] to model action-selection tasks with multiple competing goals. According to this theory, the action-selection problem in dynamic environments with competing goals can be decomposed into policy solutions for the individual goals. Consider a reaching scenario with N targets presented at different locations from the current hand state *x*_*t*_. What is the best policy that someone should follow so as to maximize the expected reward and simultaneously minimize the effort cost? According to our theory, the best policy *π*_*mix*_(*x*_*t*_) can be approximated as a weighted average of the best individual policies *π*_*j*_(*x*_*t*_) - i.e., the policies generated to reach each of the individual targets:

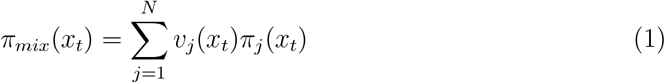

Where *v*_*j*_ is the weight of the policy *π*_*j*_ at the state *x*_*t*_ - i.e., how much this individual policy contributes to the overall policy *π*_*mix*_ followed by the agent at that state.

Each of the individual optimal policies *π*_*j*_ are generated by the minimization of the cost function in Eq.2, which describes the cost for reaching the target *j* with zero velocity, starting from the current state *x*_*t*_ and following the policy *π*_*j*_ for time instances 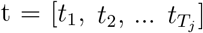

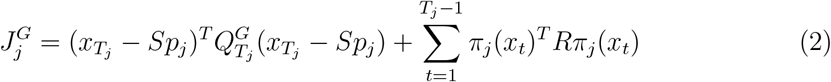

Where *T*_*j*_ is the time to contact the target *j* starting from the current state, S is a matrix that picks out the hand and the target location from the state vector *x*_*t*_. The first term of the cost function *J*_*j*_ describes the accuracy cost that penalizes policies that drive the hand away from the location of target *j*. The second term is the motor command cost that penalizes policies with high effort to reach the target *j*. Both the accuracy cost and the motor command cost characterize what we call “action cost” 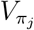 for implementing the policy *π*_*j*_ at the given state. Matrices 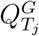 and *R* define the precision- and control-dependent costs, respectively. Given the current state *x*_*t*_ and the location of the target *j*, the controller generates the policy *π*_*j*_ that minimizes the expected cost in Eq. 2. This is a modified form of the Linear Quadratic Gaussian (LQG) regulator in optimal control theory, in which the system dynamics are linear, the cost function is quadratic and the noise is Gaussian, but with signal dependent noise [41].

#### 4.2.2 Multiple competing goals in cluttered environments

In natural tasks, we often do not act on isolated targets but in cluttered environments. To successfully reach a target in the presence of nearby obstacles, the hand should deviate away from the obstacles, in order to reach the target without collision. In the current study, we extended the computational theory presented above to model reaching movements in cluttered environments. The first step was to construct a cost function for obstacle-avoidance, similar to the one developed for reaching to targets. According to this cost, the hand should move as far away as possible from the current location of the obstacle, while exerting as little effort as possible. One of the simplest solution ideas was to use the Euclidean distance between the hand location and the location of the obstacle to define the cost function. However, this cost function is not quadratic and therefore the LQG framework is not directly applicable. Another solution, which has been extensively used by other studies, is to force the hand to pass through a via-point located nearby the obstacle [42, 43]. However, this approach biases action-selection towards particular pre-defined locations, and therefore it is not consistent with human experimental studies revealing high variability in trajectories for obstacle avoidance [7, 30].

In the current study, we proposed an alternative approach to estimate a near-optimal policy to avoid an obstacle at a given state. Instead of computing directly the policy for avoiding the obstacle, we computed the policy *π*(*x*_*t*_) to reach the obstacle from the current state without velocity constraints at the end of the movement, and then “negate” that policy. Hence, the cost function for avoiding the obstacle *k* is given as:

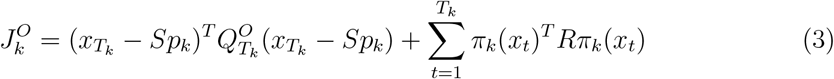

The minimization of Eq. 3 provides the optimal policy *π*_*k*_(*x*_*t*_) to reach the obstacle *k* without any velocity constraints. By negating this policy, we get a close-to-optimal policy to avoid the obstacle. The time *T*_*k*_ is the time to contact the obstacle *k*. Similar to the cost function of the targets, the first term describes the goal of the controller - i.e., arrive at the obstacle - and the second term is the effort required to arrive at the obstacle.

When multiple targets and obstacles are presented simultaneously on the field, the framework follows a mixture of policies *π*_*mix*_, which is given as a weighted average of the individual policies related to the targets (i.e., goals) *π*^*G*^(*x*_*t*_) and the obstacles *π*^*O*^(*x*_*t*_) at any given state *x*_*t*_:

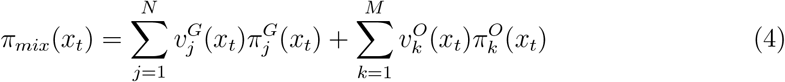

Where 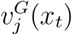 and 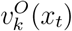 describe the contribution (i.e., weights) of each of the target-policy *j* and obstacle-policy *k*, respectively, to the mix policy *π*_*mix*_ followed by the model at the given state *x*_*t*_.

So far, we assumed that both obstacles and targets have the same influence on the mixture of the individual policies. However, experimental studies have shown individual differences in risk aversion in both humans and animals. For instance, when rushing to catch a falling object, a person might reach more directly and accept higher collision risk with nearby items, whereas a casual reach for the same object would follow a more cautious, obstacle-avoiding trajectory. To take the different levels of risk aversion into account, we introduce a scalar variable *α* that determines the risk aversion of the model. It takes values between 0 and 1, such as when *α* is close to 0, the model focuses mostly on avoiding the obstacles, and when *α* is close to 1 the model pays more attention to reaching the targets. Given the level of risk aversion, we write Eq. 4 as:

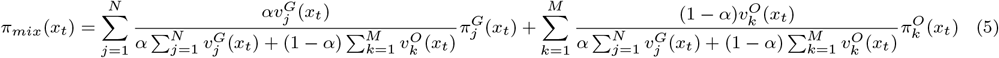

Where N and M are the number of targets and obstacles, respectively.

#### 4.2.3 Computing target and obstacle policy desirability

The remaining problem is to compute the weighting factors for the targets, *v*^*G*^(*x*_*t*_), and the obstacles, *v*^*O*^(*x*_*t*_). Consider the simplest scenario introduced in our previous work [17], in which reaches are performed in an obstacle-free environment. In this scenario, *π*_*mix*_(*x*_*t*_) is given as a weighted average of the individual policies *π*_*j*_(*x*_*t*_) related to each of the targets *j*. We called the weights *v*^*G*^(*x*_*t*_) “relative desirability” values, because they reflect how “desirable” it is to follow a particular policy *π*_*j*_(*x*_*t*_) (in this case, reach to the target *j*) with respect to the alternative options, and is given as:

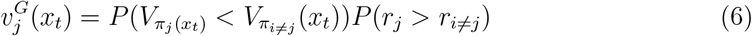

The first term is the “action-related” component of the relative desirability 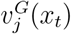 and describes the probability that reaching for the target *j* has the lowest cost (i.e., requires less effort) compared to the rest of the alternatives, at the given state *x*_*t*_. The second term is the “goods-related” component and describes the probability that selecting the target *j* will have the highest outcome (i.e., reward) compared to the alternatives, at that state. The first component of the relative desirability can be approximated by the soft-max transformation of the policy cost 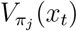 at the current state, as :

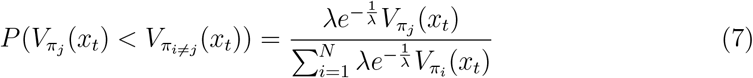

Where *λ* is the “free inverse temperature” parameter.

The second component of the relative desirability is determined by the expected benefits related to the alternative options. Our previous work [17] explored this component in simplified contexts. For instance, in a simple case where the amount of reward is fixed and equal for all goals, the relative desirability for a policy *π*_*j*_ was modeled as the probability of receiving the reward for a given target *j*:

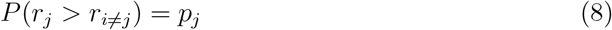

However, this model does not account for risks, such as those encountered when reaching for objects in a cluttered environment. In such scenarios, evaluating the “goods-related” component becomes more complex, as the potential reward for success must be weighed against the potential penalty for colliding with obstacles. To address this challenge, we propose a new formulation for the “goods-related” component based on the probability of collision with obstacles.

To compute this collision probability, we defined an expanded collision zone surrounding each obstacle, characterized by two concentric circles: an inner circle with diameter *d*_*inner*_ equal to the physical size of the obstacle, and an outer circle with diameter *d*_*outer*_ that delineates the extended region where collision risk is perceived. The collision cone is formed by computing tangent lines from the current hand position to both the inner and outer circles, creating a spatial region that extends from the hand toward and around the obstacle. For a given hand position at state **x**_*t*_, collision probability is computed based on its location relative to these boundaries. Points within the inner circle are assigned a collision probability of 1, while points between the inner and outer circles exhibit linearly decreasing probability:

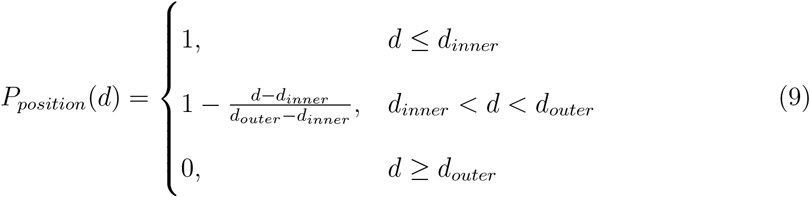

where *d* is the effective diameter of the smallest circle enclosing the hand position. To evaluate collision risk along a trajectory, we define an indicator function that incorporates both spatial location and movement direction:

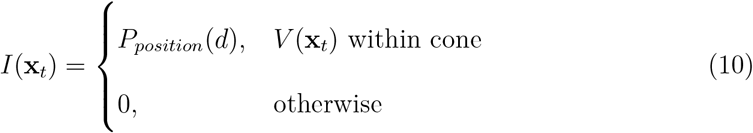

where *V* (**x**_*t*_) is the velocity vector at state **x**_*t*_. Over a trajectory spanning N future time steps, the overall collision probability is then computed as:

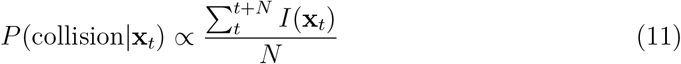

Therefore, the “goods-related” component of the relative desirability 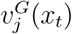 is approximated by the probability of *NOT* colliding with the obstacle (the probability of going to the target), which is defined by:

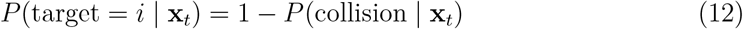

Similarly, the relative desirability for avoiding the obstacle *k*, is given as:

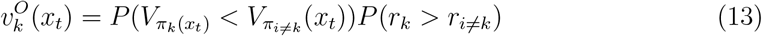

Where the first term is the action-related component of the relative desirability 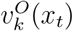, which describes the probability that avoiding the obstacle *k* has the lowest cost compared to the alternative actions, at the current state, and could be approximated by:

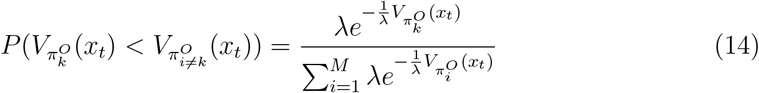

And the second term is the goods-related component of the relative desirability 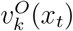 and could be simply defined by *P* (collision | **x**_*t*_).

## Acknowledgments

Research reported in this publication was supported by National Institute of Neurological Disorders and Stroke under award number U01NS132788. The content is solely the responsibility of the authors and does not necessarily represent the official views of the National Institutes of Health.

## Author Contributions

**Shan Zhong**: Methodology, Software, Formal analysis, Visualization, Writing-Original Draft, Writing-Reviewing and Editing. **Nader Pouratian**: Writing-Reviewing and Editing, Funding acquisition. **Paul Schrater**: Methodology and Editing. **Vassilios Christopoulos**: Conceptualization, Methodology, Supervision, Writing-Reviewing and Editing, Funding acquisition.

## Data and Code Availability

Data will be made available upon a reasonable request and after signing a formal data sharing agreement.

## Notes

### Competing Interest Statement

The authors have declared no competing interest.

